# No neurobehavioral evidence for reduced motivational potential of social rewards in alcohol use disorder

**DOI:** 10.1101/2025.07.17.665126

**Authors:** Yana Schwarze, Janine Stierand, Johanna Voges, Emil Zillikens, Klaus Junghanns, Oliver Voß, Frieder Michel Paulus, Sören Krach, Maurice Cabanis, Lena Rademacher

## Abstract

**Background:** The mesolimbic dopamine system plays a central role in motivating behavior. In alcohol use disorder (AUD), this system is thought to be dysfunctional, leading to hyperreactivity to alcohol-related cues. In contrast, evidence on how individuals with AUD respond to alcohol-unrelated reward cues is inconclusive, and the motivation for social rewards has not yet been investigated.

**Methods:** To address this gap, 36 individuals with AUD and 34 healthy controls performed an incentive delay task to assess social reward anticipation with a monetary and a non-reward control condition while undergoing functional magnetic resonance imaging. The ventral striatum was defined as region of interest because of its central role in neuronal circuits for motivation.

**Results:** Neither behavioral nor neuroimaging data provided any evidence of reduced motivation for social or monetary rewards in AUD. Exploratory whole-brain analyses only revealed stronger activation in the occipital/cuneal cortex in individuals with AUD than in healthy controls across all trials.

**Conclusion:** Together, these results suggest that sensitivity to social reward cues is not fundamentally impaired in AUD. Furthermore, they imply that motivational changes related to the substance do not generally alter the reward potential of alcohol-unrelated domains in AUD, opening perspectives for social-behavioral treatments for this disorder.

## 1. Introduction

The mesolimbic reward system – dopamine neurons projecting from the ventral tegmental area (VTA) to the ventral striatum – is the core of neural circuits for motivation (Stuber, 2023). Through the dopamine release, incentive salience is attributed to reward-predictive cues, making the stimulus a desirable target and initiating approach behavior (Berridge, 2012). In alcohol use disorders (AUD), this circuit is thought to be dysfunctional. According to the incentive sensitization theory (Robinson and Berridge, 2025, 1993) the dopaminergic reward system in addiction is hyperreactive to drug cues and contexts, leading to excessive “wanting” to take the drug. In line with this, functional magnetic resonance imaging (fMRI) studies examining drug cue reactivity in substance use disorders have shown aberrations in neural activation that correlate with clinical outcomes (Addiction Cue-Reactivity Initiative (ACRI) Network et al., 2024). With regard to AUD, a review by Cofresi and colleagues (2019) concluded that there is “sufficient empirical evidence to support the idea that the mechanism originally outlined in the incentive salience sensitization theory of addiction […] may be at the core of alcohol use disorder (AUD) etiology for some individuals” (pp. 912-913).

In addition to the increased incentive salience for drug-associated stimuli in substance use disorders, a reduced motivation for other, drug-unrelated rewards is frequently hypothesized, because the substance may “hijack” the neural reward system and reorganize the priority of reward processing (Tan et al., 2024; Wrase et al., 2007). A number of studies have examined the motivation for alcohol-unrelated, in most cases monetary, rewards in alcohol use disorder. For this purpose, the monetary incentive delay (MID) task (Knutson et al., 2000) is widely used. However, findings are mixed: Some studies report reduced ventral striatal activity during the anticipation of monetary rewards in AUD (Hägele et al., 2015; Romanczuk-Seiferth et al., 2015), while others found no group differences (Bjork et al., 2012; Musial et al., 2023) or even increased activity (Muench et al., 2018).

Social rewards are another type of alcohol-unrelated reward that may be of particular interest in AUD. Substance use alters social behaviors, and the mesolimbic reward system is considered to play a central part in this process (Young et al., 2011). The mechanisms are not clear, but it has been suggested that dopamine transmission in the ventral striatum mediates the rewarding aspects of social interactions and that disruptions to this process as a result of substance use affect social behaviour (Young et al., 2011). Alternatively, low social integration could result in heavier use of alcohol, a dysfunctional health behavior to cope with experienced social isolation (Havassy et al., 1991). However, while addiction neuroscience research has gained interest on social interaction and connectedness (Delgado et al., 2023), up to this date the incentive salience of social rewards per se has not yet been investigated in AUD patients. This is surprising given the importance of social factors for the treatment of addiction (Heilig et al., 2016). The present study aims to bridge this gap and investigate the motivational salience of social rewards in patients diagnosed with AUD using a well-established social adaptation (Spreckelmeyer et al., 2009) of the monetary incentive delay task by Knutson et al. (2000).

## 2. Methods

### 2.1. Sample

Forty-two individuals diagnosed with moderate or severe AUD according to DSM-5 criteria (meeting 5–11 criteria) and 38 healthy control (HC) participants were recruited between February 2019 and October 2022 (with a break between March 2020 and October 2021 due to the SARS-CoV-19 pandemic) through the Department of Psychiatry and Psychotherapy of the University of Lübeck and the AMEOS clinic in Lübeck, Germany. Healthy controls were recruited through flyers, letters, and online advertisements. All patients were in the early abstinence phase (between the 10th and 40th day) following the resolution of acute withdrawal symptoms and had no comorbid diagnosis of major depression, antisocial or borderline personality disorders, social anxiety, psychosis, or acute suicidality. Further information on inclusion criteria can be found in the Supplement. One patient and one control subject dropped out of the study. In addition, three patients and two control subjects had to be excluded from the analyses because of technical problems, excessive head motion, or sleeping. Furthermore, two patients and one control subject did not participate in magnetic resonance imaging (MRI) scanning for safety reasons and were therefore excluded from this part of the study. Thus, the final sample included 36 patients and 34 healthy controls. Both groups were matched for age and gender. Written informed consent was obtained from all participants and the study was approved by the local ethics committee of the University of Lübeck (AZ 17-077).

### 2.2. Procedure

Before study participation, all participants were screened to ensure compliance with the inclusion and exclusion criteria. Once eligible, participants attended the Center of Brain, Behavior and Metabolism in Lübeck on two separate days at the same time each day (starting at 8:30 AM ± 30 minutes). The two test days were identical in structure and included a MRI measurement, as well as numerous other measurements that were collected in the context of other research questions, such as blood samples, saliva samples, and questionnaires. The difference between the two days was that stress was induced in the participants on one day and not on the other (with the order of days counterbalanced across participants). Stress effects are not included in the present analyses and have been published elsewhere (Schwarze et al., 2024). For the present analyses, only the MRI measurements of the “control day” without stress induction are examined.

### 2.3. Social and Monetary Incentive Delay tasks

During the MRI scan, participants completed a reward paradigm that was an adaptation of the Social Incentive Delay (SID) task by Spreckelmeyer et al. (2009) and the Monetary Incentive Delay (MID) task by Knutson et al. (2000). Before the scan, the paradigm was practiced outside the MRI to ensure participants were familiar with the task. The experiment consisted of four blocks (two MID and two SID blocks) of 20 trials each, resulting in a total of 80 trials. Each trial began with a cue presentation lasting 1000 ms, indicating either a potential reward (20 trials per task, represented by a circle with a line: L) or no reward (20 trials per task, represented by an empty circle: O). This was followed by a delay period of variable length (jittered between 2025 and 3240 ms) before the target appeared. Participants were instructed to respond as quickly as possible to all target appearances, regardless of the cue type. When participants responded quickly enough in a reward trial, a picture of a smiling face (SID task) or a picture of a wallet with small amounts of money (MID task) was presented for 2500 ms. If the response was too slow, participants saw a blurred image instead. In no-reward trials, blurred images were always presented, independent of the response time. Task difficulty was standardized to a hit rate of approximately 66% by adjusting the target duration based on individual response times. If participants reacted to the flash in time, the response time window for that cue type (monetary or social, reward or no reward) was shortened by 5% and extended by 10% otherwise. The initial response time window was set to the mean response time from six practice trials conducted before the experiment plus 5%. The maximum response time was set to 500 ms for all participants. The trials in each block were pseudo-randomly arranged with inter-trial intervals jittered between 3235 and 4450 ms. MID and SID blocks were interleaved, always starting with a MID block and ending with a SID block. At the start of each block, participants were informed whether the upcoming task was MID or SID. The total duration of the paradigm was approximately 18 minutes. MRI acquisition parameters can be found in the Supplement.

### 2.4. Analysis of behavioral data

For the analysis of response times, all trials with a response time >500 ms were removed, which was the time the target was presented until the feedback was given. Then, outliers deviating more than 3 SD from the mean value of the respective condition of the respective person were excluded (0.95% of total trials), and average response times were calculated for every person and every condition. A three-way repeated-measures ANOVA was calculated with *group* (AUD vs. HC) as the between-subject factor and *task type* (social vs. monetary) and *reward* (reward vs. no reward) as within-subject factors. To correct for possible order effects of the test days, a covariate was included in the analyses, specifying whether the day without stress induction was the first or second test day. To facilitate the interpretation of non-significant results, Bayesian regression models were estimated using the brms package in R (Bürkner, 2017). Bayesian analyses were performed for the non-significant main and interaction effects observed in the ANOVA. Cauchy priors were chosen for the fixed effects, whereas the remaining priors were taken from the default_priors() function of the brms package.

### 2.5. Analysis of fMRI data

MRI data were preprocessed and analyzed using SPM12 (Wellcome Department of Imaging Neuroscience; http://www.fil.ion.ucl.ac.uk/spm) implemented in MATLAB (version R2019b; MathWorks Inc.). Functional images were slice-timed and realigned to the first image to correct for head motion. Next, they were normalized to Montreal Neurological Imaging (MNI) space using standard segmentation as implemented in SPM12 on the mean functional image and tissue probability maps. Normalization parameters were then applied to all images of the time series (resulting voxel size 3 × 3 × 3 mm). Finally, functional images were spatially smoothed with an 8 mm full width half maximum isotropic Gaussian kernel and high-pass filtered at 1/128 Hz to remove low-frequency drifts. Using a 6 mm smoothing kernel lead to no substantial changes of results (see supplementary table S1 and supplementary figures S1 and S2).

On the first level, a general linear model was created for each participant, which included four regressors with the onset times of the cues of the four conditions (reward and no reward in the social and monetary blocks) and six regressors for the onset times of feedback received. To account for motion-related artifacts, six realignment parameters and their temporal derivatives were included as nuisance regressors. Additionally, a scrubbing regressor was added, removing all volumes with a framewise displacement > 1 mm. Mean framewise displace was 0.24 ± 0.09 mm in the AUD group and 0.21 ± 0.09 mm in the HC group, and did not differ significantly between the groups (*t*(67.03) = 1.16, *p* = 0.25).

At the group level (second level), a full factorial design was set up in SPM with *group* (AUD vs. HC) as the between-subject and *task type* (social vs. monetary) and *reward* (reward vs no reward) as within-subject factors. A covariate was also set to specify whether the first or second test day was included in the analyses. In case of significant effects, post hoc t-tests were conducted to determine the direction of the observed effects. For whole-brain analyses, we used a false discovery rate (FDR)-corrected cluster-extent threshold of *p* < 0.05 based on a *p* < 0.001 voxel-level threshold as implemented in SPM12. Given the expected large effects of reward, we used a more stringent threshold of family-wise error (FWE)-corrected *p* < 0.05 for the main effect of *reward*.

Due to its central role in motivation, the ventral striatum was defined as a region of interest (ROI) for the analyses of the anticipation phase. Left and right ventral striatum masks were defined as spheres with an 8 mm radius around the peak coordinates (MNI coordinates: left, −10, 10, −2; right, 12, 14, −4) based on a meta-analysis examining reward anticipation in the ventral striatum (Diekhof et al., 2012). Mean parameter estimates were extracted from these ROIs and used in a repeated-measures ANOVA with *group* as the between-subject and *reward* and *task type* as within-subject factors. In parallel with the analysis of response times, Bayesian regression models were also estimated for the extracted parameter estimates to aid in the interpretation of non-significant results. In addition, Pearson’s correlations were computed between reward-related behavioral effects (response times in reward – no reward trials) and reward-related neural activity (parameter estimates for reward – no reward trials) in the left and right ventral striatum.

## 3. Results

### 3.1. Sample characteristics

Descriptive data on demographics, alcohol consumption, and response times in the MID and SID tasks are presented in Table 1.

**Table 1.** Sample characteristics.

|  | AUD (N = 36)<sup>a</sup> | HC (N = 34) |
| --- | --- | --- |
| Gender | 6% f, 94% m | 6% f, 94% m |
| Age (years) | 42.4 ± 8.9 | 44.0 ± 9.3 |
| Duration of abstinence (days) | 14.8 ± 6.5 | — |
| Amount alcohol (g/day) | 291.4 ± 201.7 | 2.8 ± 3.5 |
| Mean response times (ms): |  |  |
| Reward SID | 348.8 ± 36.4 | 344.3 ± 49.9 |
| Reward MID | 348.6 ± 36.5 | 342.5 ± 48.6 |
| No reward SID | 362.5 ± 40.4 | 361.4 ± 46.2 |
| No reward MID | 357.2 ± 34.8 | 354.2 ± 50.7 |

Values are given as mean and standard deviation unless specified otherwise.
<sup>a</sup>For one AUD subject, exact information about the duration of abstinence (between 10 and 40 days) and for two AUD subjects, the exact amount of alcohol before withdrawal is missing, resulting in a sample size of N=35 and N=34 for these two variables, respectively.

### 3.2. No evidence for differences between AUD and healthy controls on the behavioral level

No interaction effects of *group* with *reward* or *task type* were found for response times, providing no evidence for altered reward motivation in AUD (ANOVA *group* x *reward*: *F*(1,67) = 0.79, *p* = 0.377; *group* x *task type*: *F*(1,67) = 0.33, *p* = 0.565; *group* x *reward* x *task type: F*(1,67) = 2.67, *p* = 0.107). This was supported by a Bayesian model comparison between the full model and a reduced model that excluded the effect of interest: For all interactions involving *group*, Bayesian analyses provide anecdotal evidence against their inclusion in the model (*group* x *reward*: Bayes factor (BF_10_) = 0.58, *group* x *task type*: BF_10_ = 0.48, *group* x *reward* x *task type*: BF_10_ = 0.35). In addition, no main effect of *group* was found on response times (*F*(1,67) = 0.14, *p* = 0.71), indicating a lack of clear group differences. However, participants reacted faster to reward trials (mean: 346.1 +-42.7 ms) compared to no reward trials (mean: 358.9 +-42.9 ms; main effect of *reward* (*F*(1,67) = 50.33, *p* < 0.001). Additionally, there was a main effect of *task type* (*F*(1,67) = 5.04, *p* = 0.028), with faster response times (in reward and no reward trials) during the MID (mean: 350.7 +-42.9 ms) compared to the SID task (mean: 354.3 +-43.6 ms) and a significant *reward* x *task type* interaction (*F*(1,67) = 4.28, *p* = 0.042), reflecting a greater differentiation between reward and no reward trials in the social condition compared to the monetary condition (see Table 1, Fig. 1). To check whether these effects might be related to the order of the tasks (which consistently started with MID and ended with SID for all subjects), we repeated the analysis only for blocks two (social) and three (monetary), so that the order of the tasks is reversed. In this subanalysis, the main effect of *task type* and the *task type* x *reward* interaction were no longer significant (p > 0.502), indicating that the previously observed task effect may partially reflect an order effect.

**Figure 1.**
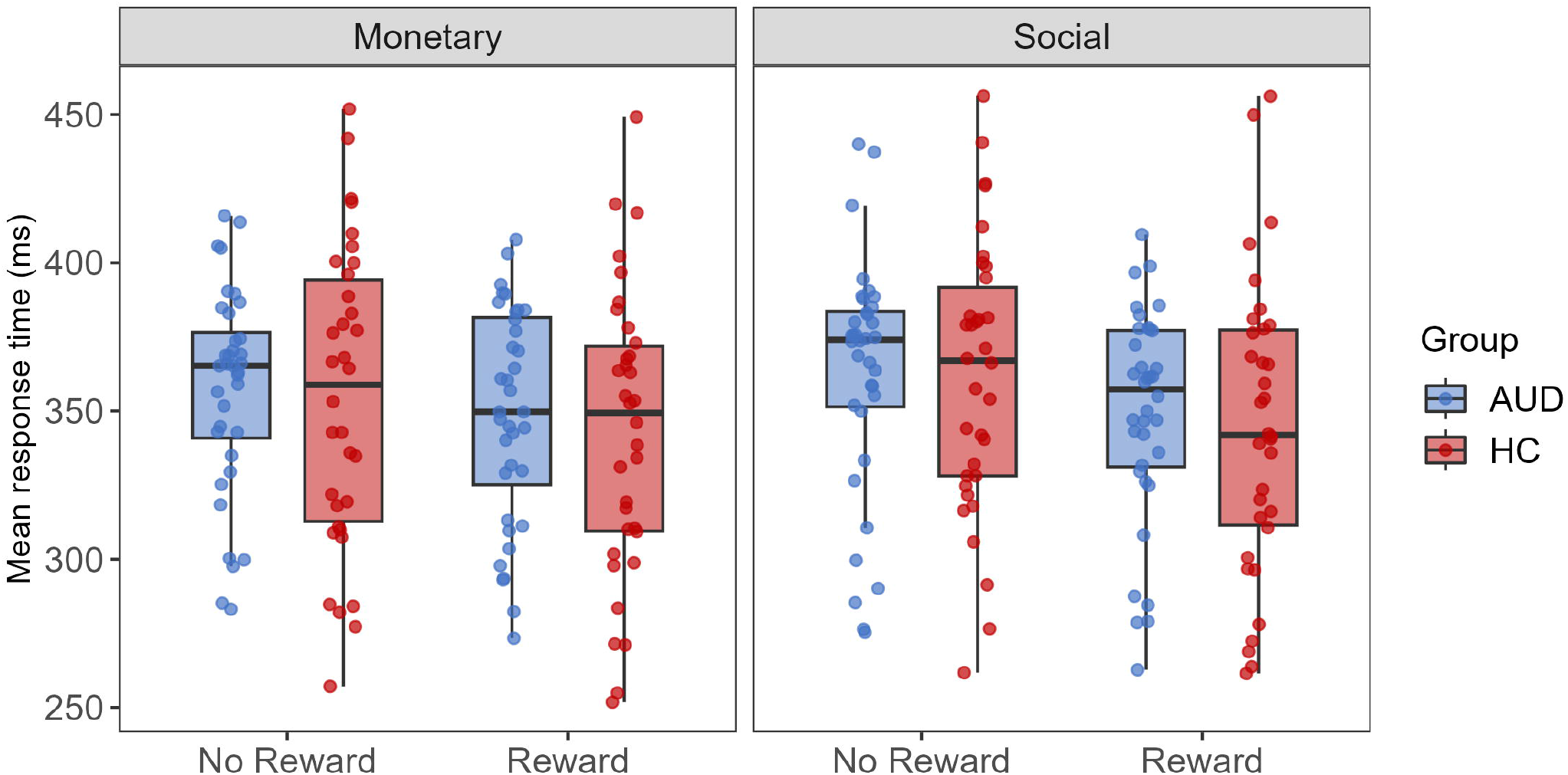
Mean response times of patients with AUD and healthy controls (HC) for reward and no reward trials of the monetary and social incentive delay task.

### 3.3. No evidence for reduced neural responses to anticipated reward in AUD

Whole brain analyses revealed no significant interaction effects between *group* and *reward* or *task type*. Significant main effects of *reward* were detected in clusters comprising regions of the striatum, occipital gyrus, supplementary motor cortex, fusiform gyrus, and insula (Table 2 and Fig. 2A). Post hoc t-tests revealed that these were all due to stronger activation during the anticipation of reward compared to no-reward trials, replicating earlier studies on reward anticipation in healthy study participants (Martins et al., 2021; Oldham et al., 2018; Wilson et al., 2018). Additionally, there was a main effect of *group* in the supracalcarine cortex/cuneus and occipital pole (see Table 2, Fig. 2B), reflecting stronger activation in patients with AUD than in healthy controls. Finally, main effects of *task type* were found in the fusiform and occipital gyrus, as well as the frontal medial and superior frontal cortex (Table 2), which reflected stronger activations during social compared to monetary reward conditions (across reward and no reward trials). As with the behavioral data, we repeated the analysis using only blocks two and three to check for an influence of order on the task effect. In this subanalysis, no significant main effect of *task type* could be found (*p* > 0.115).

**Table 2.** Activation during reward anticipation.

|  | side | MNI coordinates |  |  | Cluster size | F | p |
| --- | --- | --- | --- | --- | --- | --- | --- |
|  |  | x | y | z |  |  |  |
| **Main effect of reward** |  |  |  |  |  |  |  |
| occipital pole, lateral occipital cortex (inferior), supplementary cortex | L | -21 | -94 | -7 | 4397 | 154.96 | <0.001<sup>a</sup> |
|  |  | -27 | -88 | 8 |  | 89.68 |  |
|  |  | -6 | 5 | 50 |  | 63.41 |  |
| occipital pole, temporal occipital fusiform cortex | R | 24 | -88 | -1 | 969 | 137.49 | <0.001<sup>a</sup> |
|  |  | 30 | -46 | -19 |  | 50.99 |  |
|  |  | 36 | -58 | -13 |  | 44.65 |  |
| insular cortex, caudate, nucleus accumbens, thalamus | L/R | -30 | 23 | 2 | 1487 | 61.72 | <0.001<sup>a</sup> |
|  |  | 9 | 5 | 2 |  | 57.56 |  |
|  |  | 33 | 26 | -1 |  | 56.53 |  |
| posterior cingulate gyrus | L | -3 | -25 | 26 | 38 | 33.56 | <0.001<sup>a</sup> |
|  | L | -42 | 5 | 26 | 28 | 32.41 | 0.007 |
| **Main effect of task type** |  |  |  |  |  |  |  |
| temporal occipital fusiform cortex, occipital fusiform gyrus, lingual gyrus | L | -27 | -52 | -13 | 430 | 78.50 | <0.001 |
|  |  | -24 | -73 | -10 |  | 29.61 |  |
|  |  | -6 | -82 | -7 |  | 12.14 |  |
| temporal occipital fusiform cortex, lingual gyrus | R | 27 | -52 | -10 | 579 | 68.37 | <0.001 |
|  |  | 27 | -43 | -13 |  | 67.66 |  |
|  |  | 15 | -40 | -4 |  | 29.97 |  |
| lateral occipital cortex, cuneal cortex | L/R | 36 | -82 | 20 | 1217 | 62.81 | <0.001 |
|  |  | -33 | -88 | 17 |  | 60.04 |  |
|  |  | 0 | -88 | 23 |  | 29.96 |  |
| frontal medial cortex, frontal pole | L/R | -9 | 32 | -13 | 280 | 33.44 | 0.001 |
|  |  | 12 | 29 | -7 |  | 29.59 |  |
|  |  | 15 | 44 | -16 |  | 28.05 |  |
| superior frontal gyrus | L | -18 | 20 | 62 | 263 | 23.91 | 0.001 |
|  |  | 0 | 35 | 44 |  | 23.19 |  |
|  |  | -30 | 11 | 59 |  | 19.63 |  |
| superior frontal, middle frontal gyrus | R | 18 | 20 | 62 | 130 | 20.07 | 0.018 |
|  |  | 27 | 20 | 56 |  | 19.74 |  |
|  |  | 24 | 38 | 38 |  | 13.15 |  |
| **Main effect of group** |  |  |  |  |  |  |  |
| Supracalcarine cortex, cuneal cortex, occipital pole | R | 3 | -79 | 17 | 77 | 16.42 | 0.038 |
|  |  | 9 | -91 | 8 |  | 14.60 |  |
|  |  | 0 | -76 | 8 |  | 14.17 |  |

If not labelled otherwise, a false discovery rate-corrected cluster-extent threshold based on $p < .001$ voxel-level threshold was used.
<sup>a</sup>For these clusters, a more stringent threshold of family-wise error (FWE)-corrected $p < 0.05$ was used. MNI, Montreal Neurological Institute.

**Figure 2.**
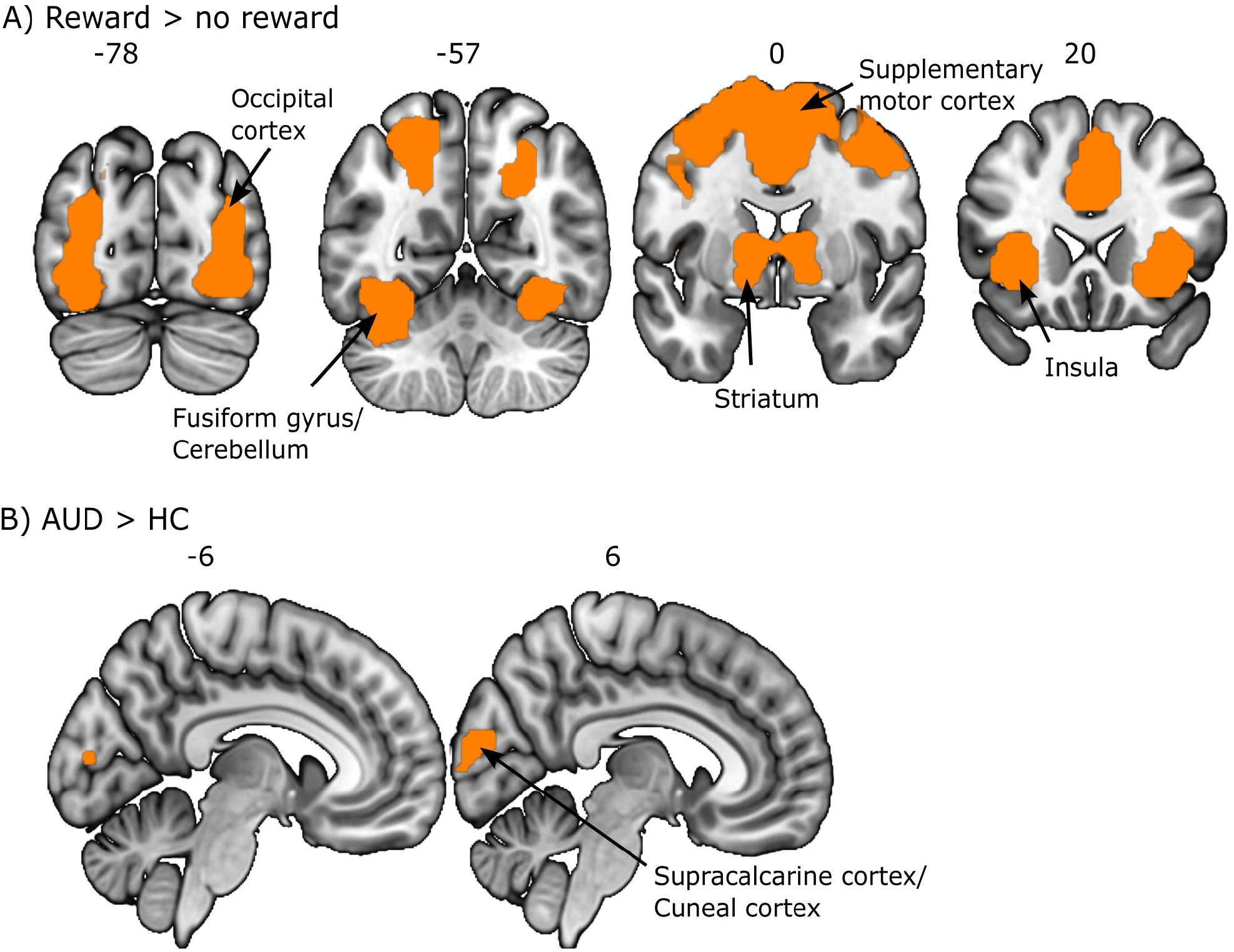
Neural activation during the anticipation phase of the Social and Monetary Incentive Delay Task. (A) Effect of reward (reward vs. no reward trials). (B) Effect of group (AUD vs. HC). A false discovery rate-corrected cluster-extent threshold of *p* < 0.05 based on *p* < .001 voxel-level threshold was used in all whole-brain analyses, except for the main effect of reward. Given the expected large effects of reward, a more stringent family-wise error-corrected threshold of *p* < 0.05 was applied.

Mean parameter estimates extracted from the left and right ventral striatum as regions of interest during reward anticipation (parameter estimates of reward trials - parameter estimates of no reward trials) negatively correlated with response times (response times of reward trials - response times of no reward trials) for both social (left: r = −0.31, p = 0.009; right: r = −0.32, p = 0.008) and monetary rewards (left: r= −0.32, p = 0.007; right: r = −0.24, p = 0.049, see supplementary figure S3), supporting the assumption that the two measures reflect motivation for rewards. The analysis of variance of parameter estimates corroborated the whole-brain findings: A main effect of *reward* (left ventral striatum: *F*(1,67) = 22.37, *p* < 0.001; right ventral striatum: *F*(1,67) = 22.42, *p* < 0.001; see Fig. 3), but no significant main effects of *task type* or *group* could be detected (p > 0.12). A significant *task type* × *reward* interaction in the bilateral ventral striatum (left: *F*(1,67) = 11.71, *p* = 0.001; right: *F*(1,67) = 10.25, *p* = 0.002) indicated a greater differentiation between reward and no reward trials in the monetary task. However, this effect was no longer present when the analysis was limited to blocks two and three (*p* > 0.686). A significant three-way interaction indicated a greater sensitivity of the left ventral striatum specifically to social rewards in the AUD compared to the HC group (*F*(1,67) = 5.40, *p* = 0.023). Group specific post-hoc comparisons showed significantly greater activation for the contrast of reward vs. no reward in both groups for monetary rewards (AUD: *p* < 0.001; HC: *p* < 0.001), while for social rewards this contrast was only statistically significant in the AUD group (AUD: *p* = 0.007, HC: *p* = 0.393). However, the groups did not significantly differ within each condition. This interaction was also no longer present when the analysis was limited to blocks two and three (*p* = 0.245). For the right ventral striatum the overall pattern was descriptively comparable but no significant three-way interaction was found (*F*(1,67) = 2.67, *p* = 0.107). No other significant interaction effects of *group* were detected. Bayesian analyses supported these findings, providing anecdotal to strong evidence against the inclusion of main effects of *group* and *task* as well as *group* x *reward* or *group* x *task type* interaction effects, with Bayes factors (BF_10_) ranging from 0.08 to 0.65 (see supplementary Table S2 for details).

**Figure 3.**
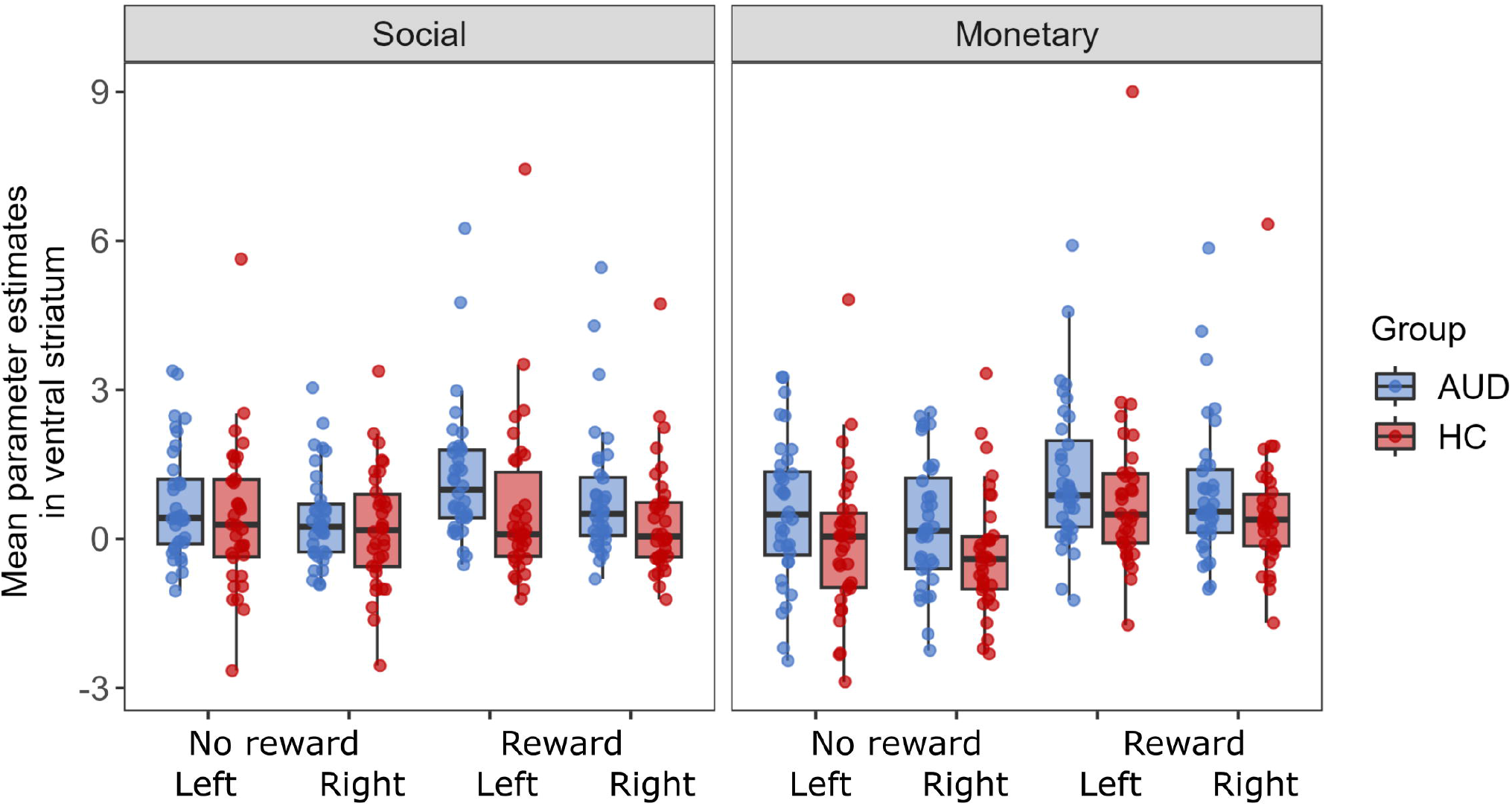
Ventral striatum response to reward cues. Average parameter estimates were extracted from the left and right ventral striatum ROI for each participant in the two groups for reward and no reward trials of each task type. AUD = Alcohol use disorder group; HC = Healthy control group.

## 4. Discussion

The present study examined the anticipation of social rewards in individuals with AUD compared with healthy control participants. Neither behavioral nor neuroimaging data provided evidence of an reduced anticipation of social reward in AUD, and similar results were observed for monetary rewards as well. These findings suggest that sensitivity to cues for social and other alcohol-unrelated rewards is not fundamentally impaired in AUD patients.

The present data showed that across both groups, response times were shorter in reward than in no reward trials. Likewise, increased neural activation during reward anticipation was observed in the striatum, occipital gyrus, supplementary motor cortex, fusiform gyrus, and insula across both groups. This finding is consistent with previous research conducted solely on healthy participants (Martins et al., 2021; Oldham et al., 2018; Wilson et al., 2018). Additionally, we found that shorter response times to potential rewards (controlled for response times in trials without rewards) were associated with increased neural activity in the ventral striatum, the core region of the neural circuits for motivation. This result is consistent with previous findings (e.g. Carruzzo et al., 2024; Kohls et al., 2013) and confirms that the anticipatory activity in the ventral striatum represents a neurobiological correlate for reward sensitivity.

The present study mainly focused on the anticipation of social rewards, which, to the best of our knowledge, has not yet been investigated in AUD. We did not find any neurobehavioral evidence of reduced motivation for social rewards in individuals with AUD. Overall, behavioral and neural data were well aligned between groups, showing the expected task effects in reaction times and responsiveness of reward related brain circuits in the ventral striatum. If at all, only the targeted analyses of fMRI parameter estimates in the left ventral striatum indicated differential responses to rewarding stimuli, however, in such a way that social stimuli showed greater rewarding potential in AUD. While this finding should not be over interpreted since it only occurred unilaterally, these data still contradict the notion of (markedly) reduced motivational potential of social rewards in AUD.

These results adds to recent findings of preserved reward learning abilities in patients with AUD and suggest that altered social behavior in AUD is not attributable to impaired reward processing per se (Jangard et al., 2025). Analogous to social rewards, we also did not find evidence of reduced motivation for monetary reward in AUD. Consistent with previous studies using the MID task, response times did not differ between patients with AUD and healthy controls (Beck et al., 2009; Bjork et al., 2012; Grodin et al., 2016; Musial et al., 2023). In line with the response times, we could not detect any evidence of altered neural processing. This finding stands in contrast to some earlier studies, including a meta-analysis of twelve studies (Zeng et al., 2023), which reported decreased ventral striatal activation during monetary reward anticipation in AUD. However, it should be noted that studies with considerable modifications to the MID (e.g., effort variations in the trials or beer and water sips instead of monetary rewards) and studies with apparently overlapping datasets were also included in the meta-analysis. Without taking these studies into account, previous research presents a heterogeneous picture of reports showing decreased, increased, or comparable ventral striatal responses in AUD compared to healthy controls. Differences in task parameters (e.g., task duration, inclusion of loss conditions, reward magnitude) and participant characteristics (e.g., shorter abstinence duration) could plausibly account for the heterogeneity of the findings.

Beyond the primary region-of-interest analysis, exploratory whole-brain comparisons revealed increased activity in AUD relative to controls across all trials (reward and no reward) in the cuneus/occipital cortex — a region for which reduced gray matter volume has been shown in AUD (Wang et al., 2018). The cuneus is implicated in shifting and sustaining attention and integrating visual information (Makino et al., 2004; Michels et al., 2008), and prior work has also linked this region to the differentiation of reward-predictive versus non-predictive cues, suggesting a role in assigning valence to potential rewards (Doñamayor et al., 2012). Additionally, increased cuneus activation in response to monetary rewards has been observed in major depressive disorder, pointing to a possible involvement of this region in reward processing across clinical populations (Zhang et al., 2013). Elevated cuneus activity may reflect altered visual attention in response to reward cues in AUD, although this interpretation remains speculative and requires further investigation.

Several limitations of the present study should be acknowledged. First, the sample predominantly consisted of male participants. Sex differences have been found in the mesolimbic dopamine release caused by alcohol (Agabio et al., 2017) and in the neural processing of rewards (Spreckelmeyer et al., 2009; Warthen et al., 2020). Therefore, the results of the present study may not be generalizable to female individuals. However, the results remain consistent after excluding the few female participants from the analyses. Second, the findings may be specific to early abstinence. Previous studies have shown that a recovery of structural, functional, and cognitive impairments can occur within the first week to month of abstinence (Parvaz et al., 2022; Powell et al., 2024). Consequently, the results may not generalise to other phases of addiction. Finally, one could argue that our study was not sufficiently powered to detect subtle group differences. While larger samples are certainly needed to explore even smaller differences between groups, it should be noted that the Bayesian analyses provided evidence supporting the absence of differential responses between groups. In addition, most previous studies reporting altered ventral striatal activity during monetary reward anticipation in AUD had smaller sample sizes (Zeng et al., 2023), rendering the current sample relatively robust in comparison.

Taken together, our results do not provide evidence that the motivation for social rewards is altered in AUD. As no group differences were found for monetary rewards either, they do not support the “hijacking” hypothesis, which states that enhanced incentive salience for alcohol-related cues in AUD comes at the expense of motivation for alcohol-unrelated rewards. Instead, our findings indicate that the fundamental processing of social and monetary reward anticipation remains intact, suggesting preserved alcohol-unrelated reward motivation in individuals with AUD. This has direct clinical relevance: Social factors play a central role in AUD treatment, and many individuals with AUD perceive alcohol as a substitute or facilitator of social reward. Accordingly, a key therapeutic goal is to help patients build alcohol-free sources of social reinforcement. Our findings suggest that this is a promising approach, given that the neural processing of other rewards — particularly social ones — remains comparable to that observed in healthy individuals, highlighting the potential of interventions that rely on the ability to respond to social rewards.

## Acknowledgements and Disclosures

We thank Alexander Schröder, Julie Forster, and Jovana Lehmann-Grube for their help in data collection.

This work was supported by the Else Kröner-Fresenius-Stiftung (to LR, Grant No. 2018_A26). JV and YS were funded by the Studienstiftung des Deutschen Volkes.

The authors declare that they have no known competing financial interests or personal relationships that could have appeared to influence the work reported in this paper.

Data and code of this study are openly available at: https://osf.io/g3ysp/

