## Supplementary material for "No neurobehavioral evidence for reduced motivational potential of social rewards in alcohol use disorder"

**Inclusion and exclusion criteria**

Participants were required to meet the following criteria (which were in part due to other research questions in the project not addressed here): age between 18 and 60 years, sufficient knowledge of the German language, scanner compatibility, no neurological disorders or brain injuries, no medication that can affect the HPA-axis or dopamine system in the past two weeks, no acute infections, no vaccination in the past three weeks, and no chronic inflammatory diseases. Additionally, patients had to meet the following criteria: diagnosis of alcohol use disorder according to the Diagnostic and Statistical Manual of Mental Disorders fifth edition (DSM-V), alcohol abstinence for at least 10 days, no comorbid diagnosis of major depression, antisocial or borderline personality disorders, social anxiety, psychosis, or acute suicidality. The inclusion criteria for control participants were as follows: no current or past psychiatric disorders, no alcohol misuse classified as ≥ 8 points in the alcohol use disorder identification test (1), no consumption of drugs (except for alcohol, nicotine, and irregular use of cannabis) in the past year, and no history of regular intake/dependency of any drug except nicotine.

**MRI acquisition**

Participants were examined in the MRI scanner at the Center of Brain, Behavior and Metabolism in Lübeck (3T Siemens MAGNETOM Skyra magnetic resonance tomography) using a 64-channel head coil. A single-shot gradient-recalled echo-planar imaging (GRE-EPI) sequence was used to acquire functional images (repetition time (TR) = 1330 ms, echo time (TE) = 25 ms, flip angle = 70°, field of view (FoV) = 560x560). Additionally, an anatomical T1 image was obtained for each participant, which was acquired using an MP-RAGE sequence (TR = 1900 ms, TE = 2.44 ms, TI = 900 ms, flip angle = 9°, resolution 1×1×1 mm3, FOV = 192×256×256 mm3; acquisition time = 4.5 minutes).

**Analysis software**

We used R version 4.3.2 (R Core Team, 2023) and the following R packages: afex v. 1.3.0 (Singmann *et al.*, 2023), arsenal v. 3.6.3 (Heinzen *et al.*, 2021), brms v. 2.22.0 (Bürkner, 2017), effectsize v. 0.8.6 (Ben-Shachar *et al.*, 2020), here v. 1.0.1 (Müller, 2020), kableExtra v. 1.4.0 (Zhu, 2024), knitr v. 1.45 (Xie, 2023), lme4 v. 1.1.35.1 (Bates *et al.*, 2015), Matrix v. 1.6.5 (Bates *et al.*, 2024), Rcpp v. 1.0.12 (Eddelbuettel and Balamuta, 2018), rstatix v. 0.7.2 (Kassambara, 2023), tidyverse v. 2.0.0 (Wickham *et al.*, 2019).

MRI data was analyzed using SPM12 (Wellcome Department of Imaging Neuroscience; http://www.fil.ion.ucl.ac.uk/spm) implemented in MATLAB (version R2019b; MathWorks Inc).

**Supplementary figures**

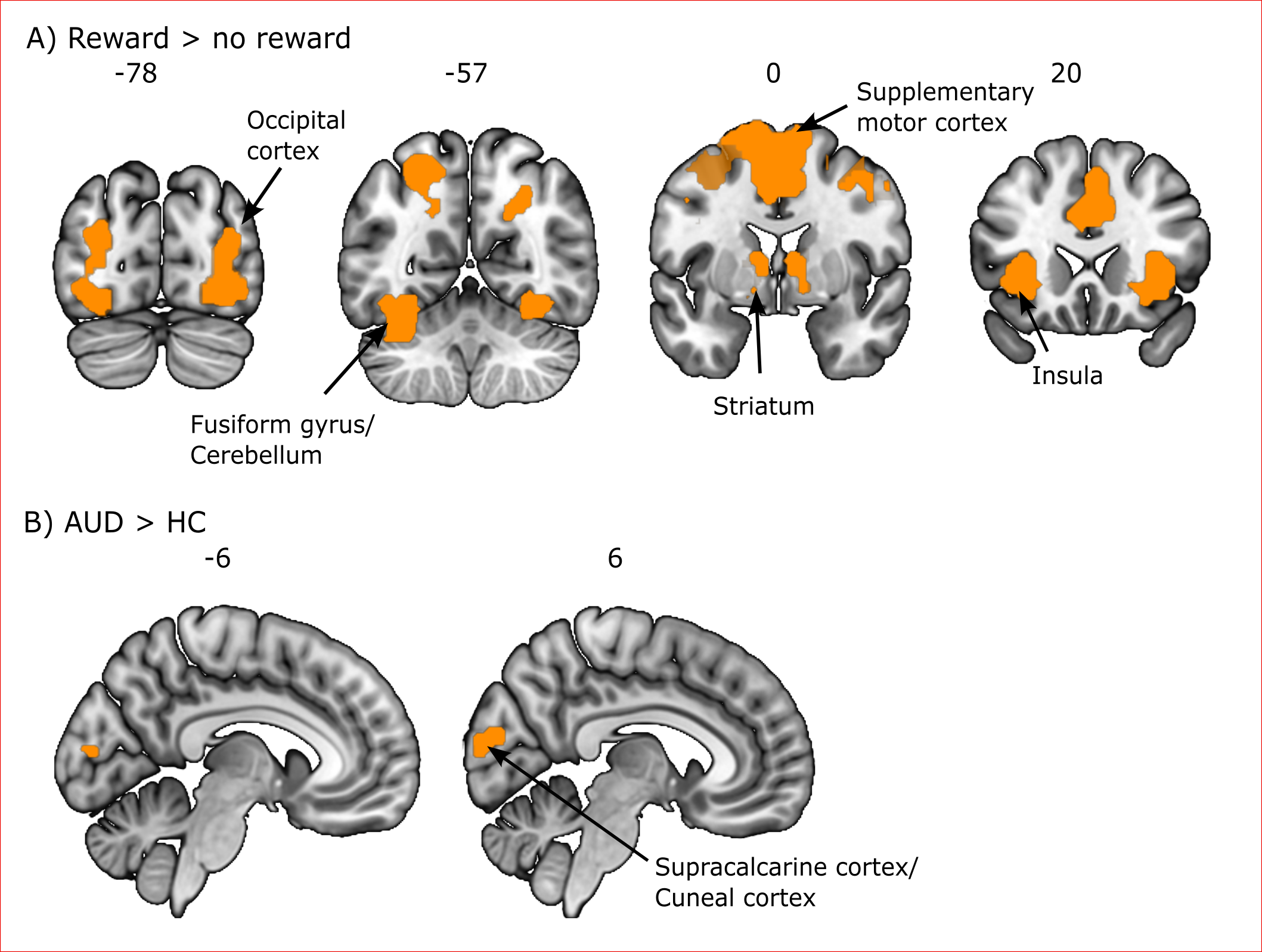

Figure S1. Neural activation during the anticipation phase of the Social and Monetary Incentive Delay Task, using a 6 mm full-width at half-maximum (FWHM) isotropic Gaussian smoothing kernel. (A) Effect of reward (reward vs. no reward trials). (B) Effect of group (AUD vs. HC). A false discovery rate-corrected cluster-extent threshold of p < 0.05 based on p < .001 voxel-level threshold was used in all whole-brain analyses, except for the main effect of reward. Given the expected large effects of reward, a more stringent family-wise error-corrected threshold of p < 0.05 was applied.

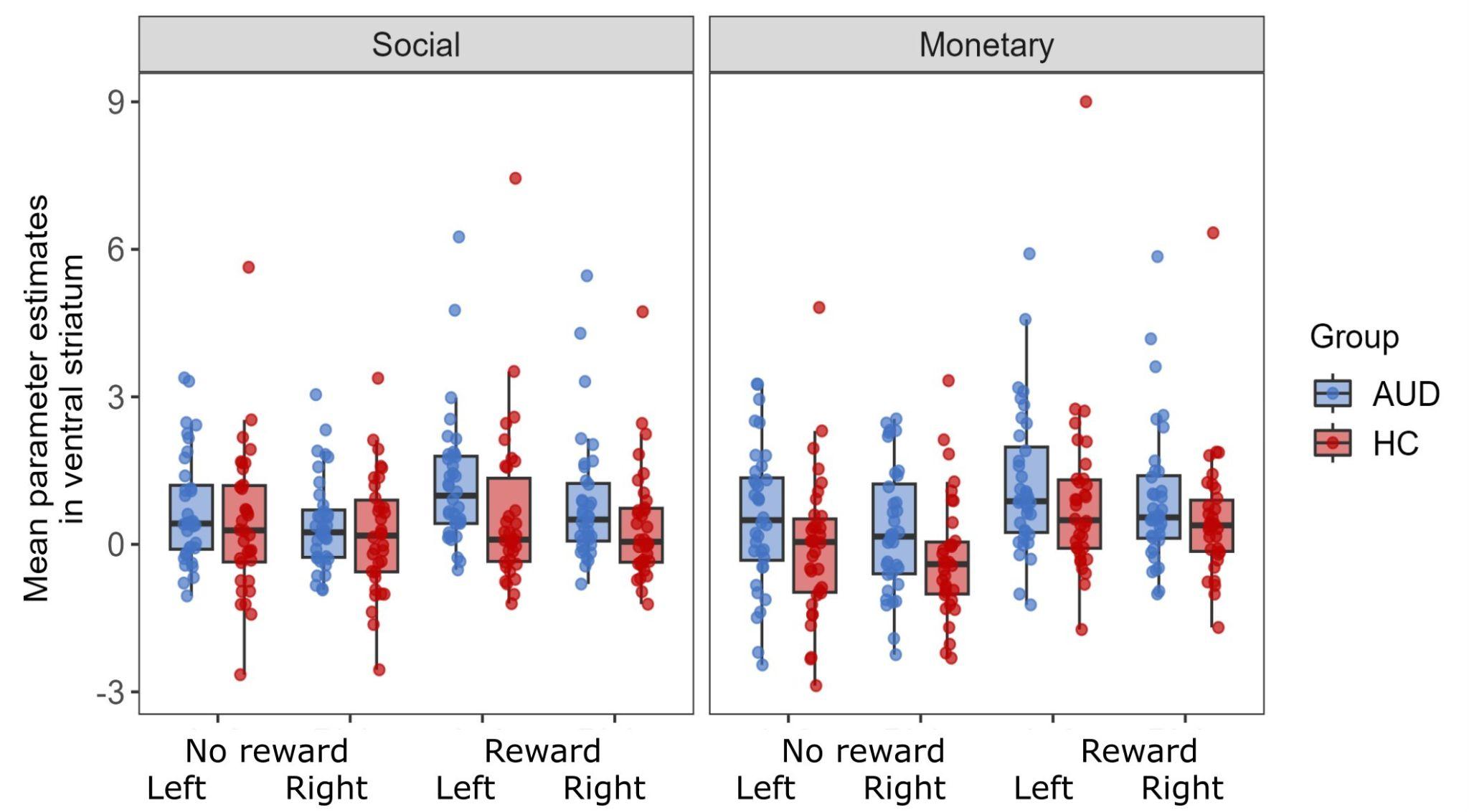

Figure S2. Ventral striatum response to reward cues, using a 6 mm full-width at half-maximum (FWHM) isotropic Gaussian smoothing kernel. Average parameter estimates were extracted from the left and right ventral striatum ROI for each participant in the two groups for reward and no reward trials of each task type. AUD = Alcohol use disorder group; HC = Healthy control group.

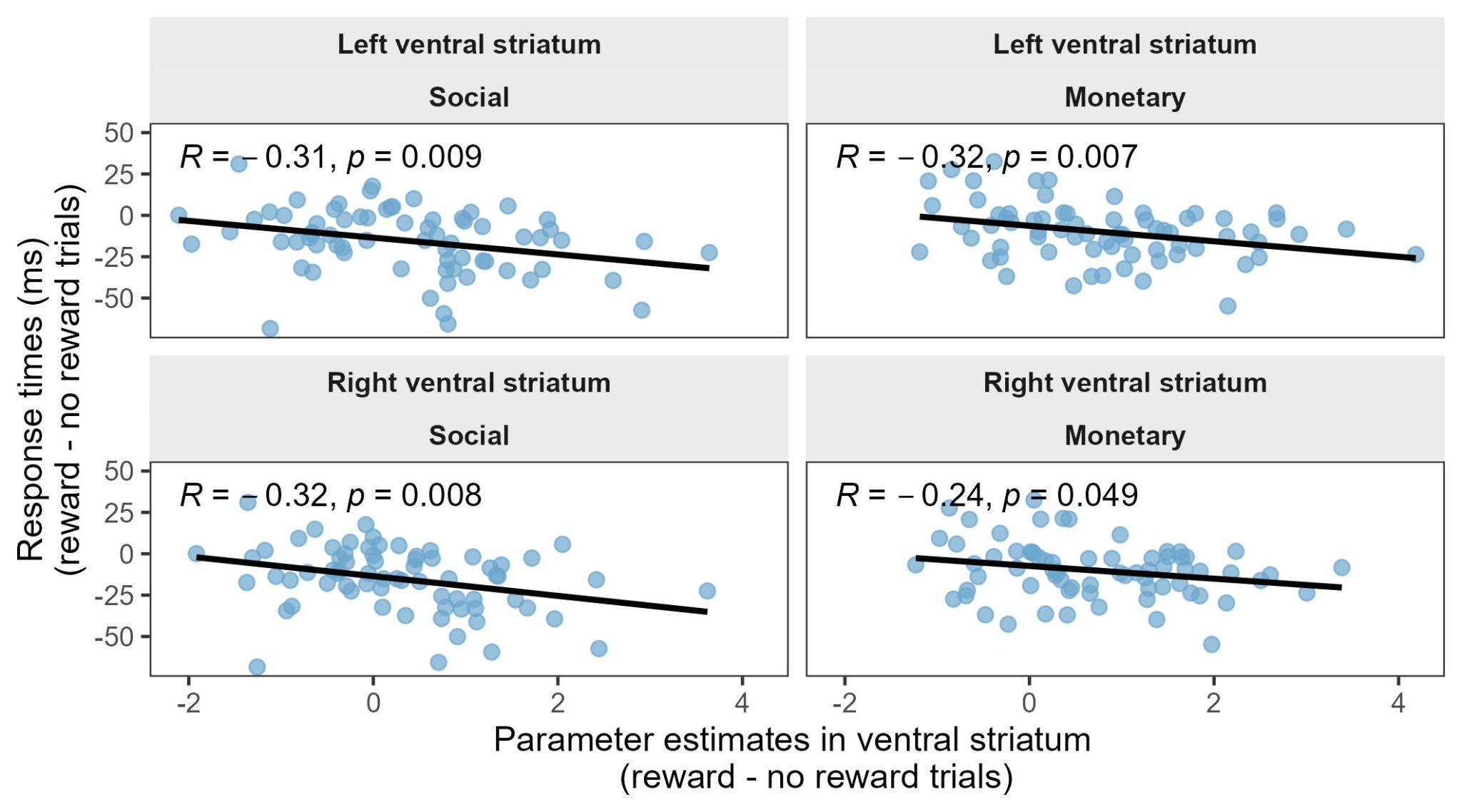

Figure S3. Correlation between neural activity in the bilateral ventral striatum during reward anticipation (parameter estimates of reward trials - parameter estimates of no reward trials) and response times (response times of reward trials - response times of no reward trials) for social and monetary rewards.

**Supplementary Tables**

Table S1. Activation during reward anticipation, using a 6 mm full-width at half-maximum (FWHM) isotropic Gaussian smoothing kernel.

|  | side | MNI coordinates | | | Cluster size | F | p |
| --- | --- | --- | --- | --- | --- | --- | --- |
|  |  | x | y | z |  |  |  |
| **Main effect of reward** |  |  |  |  |  |  |  |
| occipital pole, lateral occipital cortex (inferior), temporal occipital fusiform cortex | L | -24 | -91 | -7 | 1101 | 146.22 | <0.001^a^ |
|  |  | -27 | -88 | 8 |  | 88.20 |  |
|  |  | -33 | -55 | -16 |  | 61.61 |  |
| occipital pole, temporal occipital fusiform cortex | R | 24 | -91 | -4 | 684 | 126.43 | <0.001^a^ |
|  |  | 30 | -43 | -19 |  | 49.90 |  |
|  |  | 27 | -61 | 38 |  | 45.12 |  |
| insular cortex | L | -30 | 23 | 2 | 156 | 63.51 | <0.001^a^ |
| juxtrapositional lobule cortex, paracingulate gyrus | L/R | -3 | 8 | 53 | 1384 | 60.55 | <0.001^a^ |
|  |  | -3 | -4 | 56 |  | 51.43 |  |
|  |  | 6 | 8 | 50 |  | 50.96 |  |
| caudate, insula, thalamus, putamen | L/R | 9 | 5 | 2 | 710 | 59.04 | <0.001^a^ |
|  |  | 33 | 23 | 2 |  | 56.18 |  |
|  |  | -9 | 8 | 2 |  | 54.88 |  |
| precentral gyrus | R | 54 | 5 | 44 | 150 | 43.20 | <0.001^a^ |
|  |  | 39 | -4 | 50 |  | 36.52 |  |
|  |  | 33 | -1 | 44 |  | 33.27 |  |
| precentral gyrus | L | -39 | 5 | 26 | 17 | 34.19 | <0.001^a^ |
| cingulate gyrus | L | -3 | -25 | 26 | 17 | 33.97 | <0.001^a^ |
| precentral gyrus | R | 45 | 8 | 29 | 22 | 30.99 | <0.001^a^ |
| **Main effect of task type** |  |  |  |  |  |  |  |
| temporal occipital fusiform cortex, occipital fusiform gyrus, parahippocampal gyrus | L | -27 | -49 | -13 | 345 | 74.43 | <0.001 |
|  |  | -24 | -73 | -10 |  | 26.00 |  |
|  |  | -15 | -37 | -10 |  | 13.33 |  |
| temporal occipital fusiform cortex, occipital fusiform gyrus, lingual gyrus | R | 27 | –58 | -7 | 466 | 70.50 | <0.001 |
|  |  | 27 | -43 | -13 |  | 61.13 |  |
|  |  | 24 | -70 | -13 |  | 35.75 |  |
| lateral occipital cortex | R | 36 | -82 | 23 | 260 | 54.06 | <0.001 |
|  |  | 39 | -82 | 14 |  | 50.13 |  |
|  |  | 12 | -82 | 41 |  | 17.06 |  |
| lateral occipital cortex | L | -33 | -88 | 17 | 516 | 52.53 | <0.001 |
|  |  | -27 | -88 | 23 |  | 47.15 |  |
|  |  | -27 | -79 | 14 |  | 35.82 |  |
| paracingulate gyrus, subcallosal cortex | L/R | -6 | 32 | -13 | 99 | 29.59 | 0.002 |
|  |  | 12 | 29 | -7 |  | 25.64 |  |
|  |  | 3 | 29 | -16 |  | 25.12 |  |
| superior frontal gyrus, paracingulate gyrus | R | 0 | 35 | 44 | 68 | 25.77 | 0.009 |
|  |  | 12 | 29 | 38 |  | 18.82 |  |
|  |  | 0 | 41 | 32 |  | 12.31 |  |
| superior temporal gyrus, temporal pole | R | 57 | -1 | -7 | 51 | 25.46 | 0.027 |
|  |  | 51 | 8 | -13 |  | 23.61 |  |
|  |  | 51 | -7 | -1 |  | 13.54 |  |
| putamen | L | -27 | 5 | -1 | 49 | 22.73 | 0.029 |
|  |  | -21 | 2 | 14 |  | 15.79 |  |
| superior frontal gyrus, middle frontal gyrus | L | -12 | 23 | 62 | 98 | 22.09 | 0.002 |
|  |  | -30 | 11 | 59 |  | 20.67 |  |
|  |  | -21 | 23 | 53 |  | 17.23 |  |
| superior frontal gyrus | R | 27 | 23 | 56 | 75 | 18.94 | 0.006 |
|  |  | 15 | 17 | 62 |  | 17.77 |  |
|  |  | 9 | 26 | 59 |  | 14.08 |  |
| **Main effect of group** |  |  |  |  |  |  |  |
| Supracalcarine cortex, lingual gyrus, cuneal cortex, occipital pole | R | 3 | -79 | 17 | 80 | 20.32 | 0.004 |
|  |  | 0 | -73 | 8 |  | 18.19 |  |
|  |  | 6 | -88 | 14 |  | 14.36 |  |

If not labelled otherwise, a false discovery rate–corrected cluster-extent threshold based on *p* < .001 voxel-level threshold was used.

^a^For these clusters, a more stringent threshold of family-wise error (FWE)-corrected p < 0.05 was used.

MNI, Montreal Neurological Institute.

**Table S2. Bayes factors (BF_10_) of extracted parameter estimates of left and right ventral striatum.**

|  | **left ventral striatum** | **right ventral striatum** |
| --- | --- | --- |
| Group | 0.648 | 0.573 |
| Task type | 0.179 | 0.126 |
| Group x reward | 0.117 | 0.084 |
| Group x task type | 0.085 | 0.112 |

**Table S3: Model fit of brms models**.

| **Dependent variable** | **Model fit** | | | |
| --- | --- | --- | --- | --- |
|  | **max. R-hat** | **min. Bulk-ESS** | **min. Tail-ESS** | **R^2^** |
| Response times | 1.00 | 1235 | 2963 | 0.92 |
| Parameter estimates left striatum | 1.00 | 1607 | 2391 | 0.86 |
| Parameter estimates right striatum | 1.00 | 1636 | 2408 | 0.81 |

R-hat values were close to 1.00 for all parameters, indicating good convergence (R-hat ≈ 1); maximum R-hat per model is reported. The minimum Bulk-ESS and Tail-ESS values across all parameters per model are reported as measures of sampling efficiency for means and tails (higher values indicate better efficiency). Bayesian R² summarizes model fit as the proportion of variance explained by the model.

Formula: DV ~ group_z + task_type_z + reward_z + group_z:task_type_z + group_z:reward_z + task_type_z:reward_z + group_z:reward_z:task_type_z + on_which_day_TSST_z + (1 + task_type_z + reward_z | subj_num)
